# Interactions with stromal cells promote a more oxidized cancer cell redox state in pancreatic tumors

**DOI:** 10.1101/2020.10.20.347658

**Authors:** Rupsa Datta, Allison N. Lau, Sharanya Sivanand, Logan Florek, Jeffrey Wyckoff, Melissa C. Skala, Matthew G. Vander Heiden

## Abstract

Access to electron acceptors supports oxidized biomass synthesis and can be limiting for cancer cell proliferation, but how cancer cells overcome this limitation in tumors is incompletely understood. Non-transformed cells in tumors can help cancer cells overcome metabolic limitations, particularly in pancreatic cancer, where pancreatic stellate cells (PSCs) promote cancer cell proliferation and tumor growth. However, whether PSCs affect the redox state of cancer cells is not known. By taking advantage of the endogenous fluorescence properties of reduced nicotinamide adenine dinucleotide cofactors and oxidized flavin adenine dinucleotide, we use optical imaging to assess the redox state of pancreatic cancer cells and PSCs and find that the redox state of cancer cells is more reduced while the redox state of PSCs is more oxidized. Direct interactions between PSCs and cancer cells promote a more oxidized state in cancer cells, suggesting that metabolic interactions between cancer cells and PSCs is a mechanism to overcome the redox limitations of cell proliferation in pancreatic cancer.

## Introduction

Tumors are a heterogeneous collection of cell types, and cancer cells must adapt metabolism to use available nutrients and proliferate^1^. A prominent feature of pancreatic ductal adenocarcinoma (PDAC) is that cancer cells are a relatively rare cell type in tumors, with interactions between cancer cells and non-cancer cells affecting many properties of this cancer including cancer progression and drug sensitivity^2–6^. Diverse populations of stromal and immune cells within the tumor microenvironment can also influence nutrient utilization, and there is evidence for metabolic competition and cooperation between cancer cells and other cell types in pancreatic cancer^2,3,5,7–9^. Efforts to target the desmoplastic stroma, including pancreatic stellate cells (PSCs), have been challenging because stromal cells can both promote and restrain tumor progression^10–14^. Thus, identifying specific ways that stromal cells such as PSCs interact with cancer cells to promote tumor growth is necessary to target the stroma and improve pancreatic cancer treatment.

Oncogenic activation of KRAS is an early event in pancreatic tumor progression and supports metabolic reprogramming of cancer cells as well as the retention of stromal cells within the tumor^15^. Moreover, activation of PSCs, a major resident cell type within the pancreas, is observed prior to tumor formation, suggesting that cooperative interactions between PSCs and cancer cells influence early disease progression^16^. Cancer associated fibroblasts (CAFs), including activated PSCs, can also affect therapy response. CAFs and associated desmoplasia can account for up to 90% of the tumor, leading to a high interstitial pressure that limits drug delivery^17^. CAFs can also produce metabolites that compete for uptake of nucleoside analog chemotherapy^2^. As a result, the presence of CAFs, including PSCs, can promote pancreatic tumor growth and impact response to therapy^9,18,19^.

Pancreatic cancer organoids derived from genetically engineered mouse models mimic some aspects of human PDAC and can be used to study tumor and stromal cell interactions in a co-culture setting where morphology based identification of cancer cells and PSCs is feasible^4,9,20,21^. Another advantage of pancreatic cancer organoid co-cultures is that metabolic interactions between two defined cell types can be studied in an *in vitro* setting that is more tractable for metabolic studies than analysis of whole tumors^4,9,22^. PSCs can support the growth of tumor organoids in minimal media conditions that are not conducive to culture of organoids derived from only cancer cells^4,9^. This argues that co-culture of cancer cells and PSCs in organoid cultures is a system where metabolic interactions between cell types are active. However, exactly which media conditions can be replaced by PSCs to support cancer cell proliferation are not known.

Access to electron acceptors to support regeneration of oxidized nicotinamide adenine dinucleotide (NAD^+^) as a cofactor for oxidized biomass synthesis can be limiting for cancer cell proliferation^23–31^. This problem is particularly apparent in PDAC tumors, where levels of oxygen, an important biological electron acceptor, are extremely low^32^. Thus, how pancreatic cancer cells dispose of excess electrons to regenerate NAD^+^ is unknown, and one hypothesis is that interactions with PSCs allow cancer cells to overcome this metabolic limitation of tumor growth.

The rapid time scales of metabolic processes can be an impediment to studying metabolic interactions between cell types in a mixed population. The time needed to dissociate either organoids or tumors and isolate individual cell types is long relative to the turnover of many metabolites, and dissociating and sorting cells can affect metabolic pathways known to be responsive to cell stress^9,33–35^. One approach that circumvents these limitations is imaging metabolic changes in intact systems such as organoids and tumors. A parameter that is amenable to imaging with minimal perturbation of cell-cell interactions is assessment of cell redox state, which can be assessed at the single cell level even when multiple cells types are present. An optical redox ratio can be measured using label-free, high-resolution multiphoton microscopy of redox co-factor autofluorescence^36–39^. This technique takes advantage of the fact that only the reduced form of nicotinamide adenine dinucleotide (NADH) and nicotinamide adenine dinucleotide phosphate (NADPH) and the oxidized form of flavin adenine dinucleotide (FAD) are fluorescent.

NADPH has overlapping spectral properties with NADH, hence the autofluorescence measured is derived from both NADH and NADPH. However, while the NADH/NAD^+^ ratio is less oxidized in cells than the NADPH/NADP^+^ ratio, these ratios are coupled, suggesting that changes in NAD(P)H fluorescence can be a surrogate for changes in the reduced state of these cofactors^40–44^. Likewise, the FAD/FADH2 ratio is coupled to redox reactions in pathways that also involve NAD(P)H, enabling changes in FAD fluorescence to reflect changes in oxidized cofactors in oxidation-reduction reactions^37,45,46^. Thus, the optical redox ratio, defined as the ratio of fluorescence intensity of NAD(P)H to FAD, provides a quantitative measurement of the relative oxidation-reduction state of individual cells^36,38,47^. This optical redox measurement has been used to study the metabolic state of myriad cell types including cell populations found in tumors such as cancer cells^18–50^ and immune cells^51–53^. Multiphoton microscopy also allows optical sectioning to accommodate metabolic imaging of 3D biological samples like organoids and intravital imaging of mouse tumors ^36,54^ Finally, the non-destructive nature of the technique allows for measurements of redox state over time at a single cell level without needing to dissociate cells.

In this study, we use optical redox imaging to show that PSCs can impact the redox state of pancreatic cancer cells in organoid culture. PDAC cancer cells are more reduced relative to PSCs, but physical interactions between cell types promote a more oxidized state in cancer cells. These data argue that a metabolic symbiosis exists whereby fibroblasts facilitate oxidation reactions in cancer cells to support proliferation even in the absence of exogenous electron acceptors.

## Results

### Co-culture with PSCs supports pancreatic cancer cell proliferation in the absence of exogenous pyruvate

Prior studies have found that PSCs can secrete metabolites and other factors that impact pancreatic cancer cell metabolism and proliferation^2,7,55^, and PSCs can support pancreatic cancer cell proliferation in organoid culture^2,4,7,9,55^. To further examine how PSCs might support pancreatic cancer cell metabolism and enable organoid growth, we co-cultured primary or immortalized murine PSCs^4,9^ with PDAC cells from tumors arising in the *LSL-Kras^G12D^; Pdx1-Cre; Trp53^fl/fl^* (KP^-/-^C) autochthonous mouse PDAC model^56^ as 3D organoids. Because prior work suggested that co-culture with PSCs can enable organoid growth in minimal media^4,9^, we first examined whether contact between PSCs and cancer cells was necessary for this phenotype using a transwell system. In this experiment, media is shared between PDAC cancer cells grown as 3D organoids and 2D cells grown on the transwell inserts, but organoids are not in physical contact with PSCs or other cell types included in the transwell system. Specifically, PDAC cells cultured as 3D organoids in minimal media were seeded in the bottom chamber in Matrigel domes, and murine PSCs, mouse embryonic fibroblasts (MEFs), or PDAC cells were seeded in the upper chamber of the transwell system (Supplementary Figure 1). We found that adding MEFs or PSCs in the upper chamber of the transwells increased the number of organoids formed in the lower chamber when compared with the number of organoids observed when cultured without cells added to the upper chamber of the transwell, or when PDAC cells were seeded in the upper chamber (Figure 1a). This increase in organoid number was similar when either primary or immortalized PSCs were added to the upper well of the transwell. Interestingly, although culturing PDAC cell organoids with any other cell type added to the upper chamber of the transwells increased the size of a subset of the organoids that formed (Figure 1b), organoid number was specifically enhanced when cultured with PSCs (Figure 1a). These results are in agreement with previous observations that both co-cultures and transwell cultures involving PSCs and PDAC cells can increase cancer cell proliferation^4,7,9^. Moreover, these data argue that interactions with other cell types can affect organoid size and number differently.

**Figure 1.**
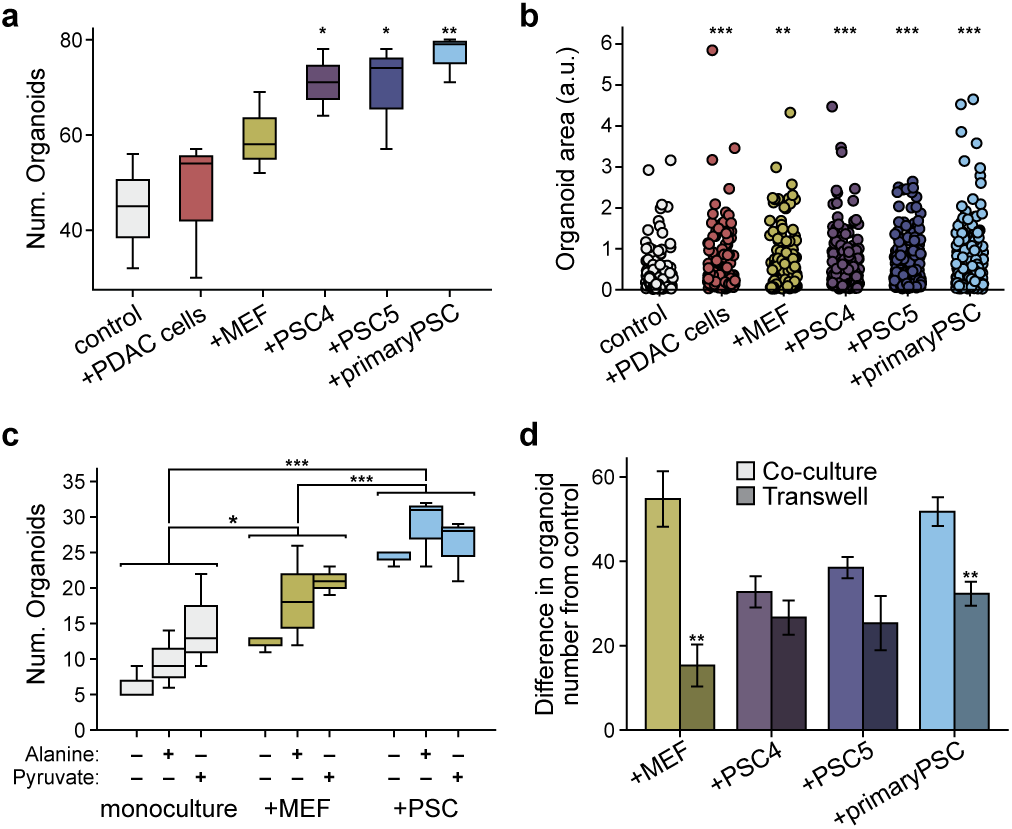
Pancreatic stellate cells support pancreatic cancer cell organoid growth. (**a-b)** Murine pancreatic ductal adenocarcinoma (PDAC) cells were cultured as 3D organoids in Matrigel domes in the bottom chamber of transwell plates alone (control), or with murine PDAC cells (+PDAC cells), mouse embryonic fibroblasts (+MEF), immortalized PSCs (+PSC4, +PSC5), or primary pancreatic stellate cells (+primary PSC) in standard 2D culture in the upper chamber of the transwell. The number of organoids (Num. Organoids) formed (**a**) and the relative area of the organoids (**b**) was quantified from brightfield images. Three independent wells were assessed per condition and statistical significance for any differences from control was determined using one-way analysis of variance (ANOVA) with posthoc Dunnett’s test. At least 3 images were analyzed per condition. (*** p< 0.001; ** p< 0.01; * p< 0.05). Error bars represent the 95% confidence interval. (**c**) PDAC cells cultured as 3D organoids were cultured alone (monoculture) or co-cultured with MEFs (+MEF) or immortalized PSCs (+PSC) included within the Matrigel domes. For each condition, cells were exposed to DMEM without pyruvate and supplemented with dialyzed serum as well as 1 mM alanine or 1 mM pyruvate as indicated. The number of organoids (Num. Organoids) formed was quantified from brightfield images. Three independent wells were assessed per condition, and any significant differences were determined using one-way ANOVA with posthoc Tukey’s test. (*** p< 0.001; ** p< 0.01; * p< 0.05). Error bars represent the 95% confidence interval. (**d**) Difference in organoid number between PDAC cells cultured as 3D organoids in co-culture with MEF, PSC4, PSC5, or primary PSC compared to PDAC cells cultured as 3D organoids alone (control) (light colored bars, calculated from data presented in Supplementary Figure 2f). Also plotted are the difference in organoid number for PDAC cells cultured as 3D organoids in a transwell bottom chamber with MEFs, PSC4, PSC5 or primary PSC in the upper chamber compared to PDAC cells cultured as 3D organoids without any cells in the upper chamber (control) (darker colored bars, calculated from data presented in (a)). For all conditions, the cultures were grown in media without exogenous pyruvate or alanine. n= 4 images were analyzed for co-cultures and n = 3 images were analyzed for transwells for each condition. Significant differences in organoid number from co-culture conditions compared to transwell conditions were assessed using T-test is indicated (*** p< 0.001; ** p< 0.01; * p< 0.05). Error bars represent standard deviation.

We next explored which metabolites in PSC- and cancer cell-conditioned media could be promoting the increase in pancreatic cancer organoid number. Since alanine has been shown to be secreted by PSCs^7^, and pyruvate has been shown to promote proliferation of multiple cancer cells types including pancreatic cancer^23,24,57,58^, we first examined whether the organoid growth-promoting role of PSCs could be replaced by addition of exogenous pyruvate or alanine to the media of organoid monocultures (Supplementary Figure 1). Strikingly, when grown in monoculture, pancreatic cancer cell organoid number was approximately doubled in the presence of exogenous pyruvate (Figure 1c, Supplementary Figure 2a-c). A possible trend toward a slight increase in organoid size was observed when either exogenous alanine or pyruvate is added to cancer cell organoid monocultures, although this difference was not significant (Supplementary Figure 2d). These results indicate that the presence of exogenous pyruvate can promote increased numbers of pancreatic cancer-derived organoids, and that including pyruvate in the medium can replace access to PSCs to increase the number of organoids.

To examine whether the growth-promoting effects of stromal cells supersedes the effects of exogenous pyruvate or alanine, PDAC cells grown as organoids were co-cultured with immortalized PSCs or MEFs (Supplementary Figure 1), with and without addition of these exogenous metabolites (Figure 1c, Supplementary Figure 2a-d). Compared to organoid monocultures containing only PDAC cells that were supplemented with pyruvate or alanine, we observed a further increase in the total number of organoids when co-cultured with MEFs (p<0.05), and this effect was further increased when the PDAC cells were co-cultured with PSCs (p< 0.001) (Figure 1c, Supplementary Figure 2c). Of note, this affect was observed regardless of whether alanine or pyruvate were supplemented in the media, and PSCs were significantly better at enhancing the number of organoids formed compared to MEFs (p<0.001) (Figure 1c, Supplementary Figure 2b). Co-culture with MEFs and PSCs may also cause a slight increase in the size of some organoids compared to monoculture, however the overall difference in size observed was not significant (Supplementary Figure 2d). Of note, exogenous pyruvate has a greater effect on organoid number in monoculture than it does in co-culture with PSCs (Figure 1c, Supplementary Figure 2a), suggesting that exogenous pyruvate provides less of an added growth benefit in the presence of PSCs. Similarly, organoid number was increased when organoids were in co-culture with stromal cells to a greater degree than that observed when pyruvate is added to monocultures (Figure 1c, Supplementary Figure 2e). Notably, co-culture with MEFs or PSCs increased organoid size more than was observed following pyruvate addition to monocultures, or when including stromal cells in transwell cultures (Figure 1d, Supplementary Figure 2f). Taken together, these data argue that interactions between cancer cells and stromal cells can reduce the requirement for pyruvate or alanine addition to the media. Furthermore, physical contact between cell types may contribute to this effect, and physical interactions mitigate the need for pyruvate more than for alanine.

### Optical redox imaging reveals redox difference between cell types in 2D monoculture as well as in tumor-PSC co-culture

Both pyruvate and alanine can contribute carbon to central carbon metabolism, but unlike alanine, which has a redox state that is equivalent to lactate^59^, exogenous pyruvate can serve as an electron acceptor to support biomass production, which can be limiting for proliferation in some contexts^23,24,26,57^. To study whether metabolic interactions between PDAC cells and PSCs overcome a requirement for pyruvate as an electron acceptor to support cancer cell proliferation, we first asked whether PDAC cells and PSCs exhibit a redox state difference when cultured individually in standard cell culture conditions. We measured the NADH/NAD^+^ ratio in murine PDAC cells derived from the KP^-/-^C mouse PDAC model and in immortalized mouse PSCs cultured in media with or without pyruvate (Figure 2a). PSCs had a more oxidized NADH/NAD^+^ ratio compared to cancer cells irrespective of the presence of pyruvate in the growth media, and unlike PDAC cells, their NADH/NAD^+^ ratio was not affected by addition of pyruvate to the media (Figure 2a). These data suggest that PDAC cells and PSCs exhibit different redox states when cultured separately in standard 2D conditions.

**Figure 2.**
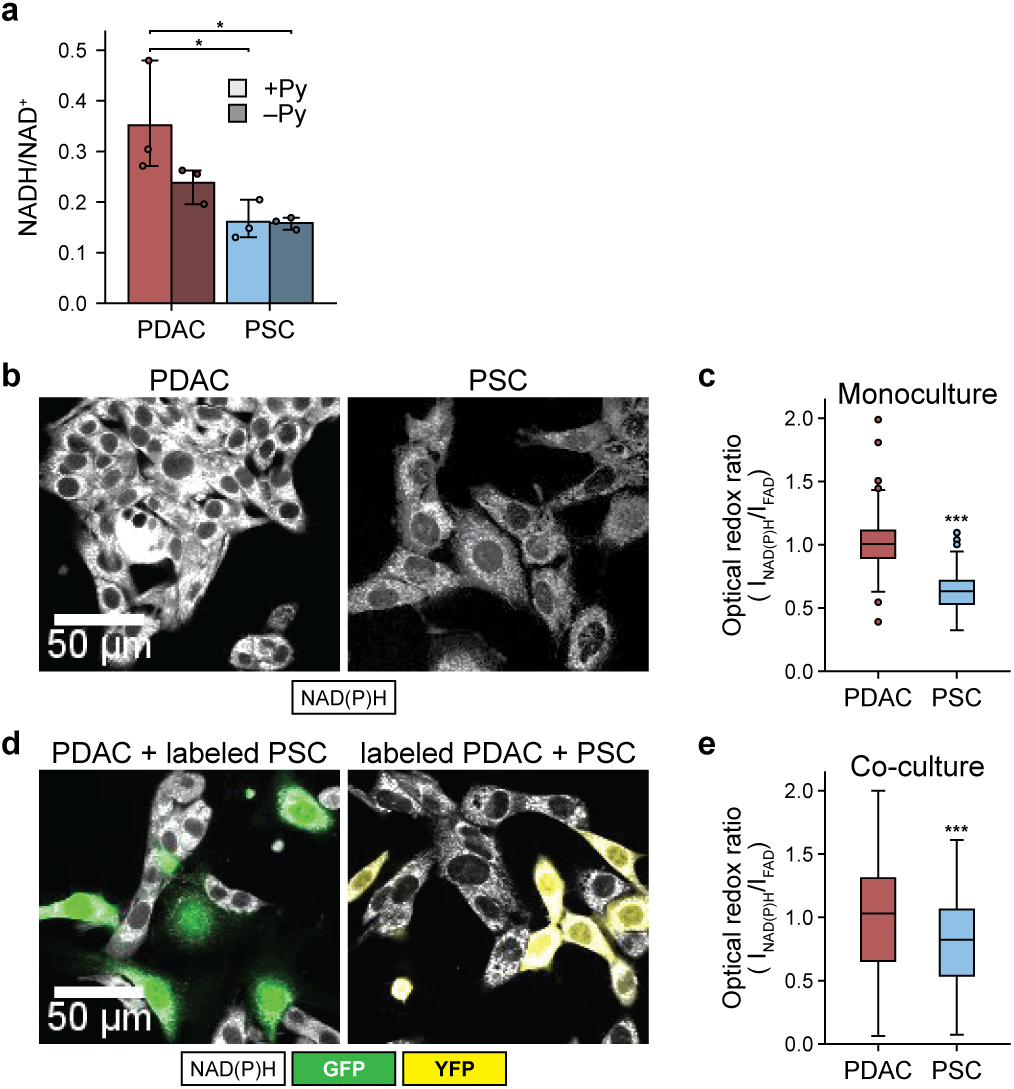
PDAC cells and PSCs in 2D culture exhibit different redox states. (**a**) Biochemical analysis of NADH/NAD^+^ ratio in PDAC cells isolated from *LSL-KRas^G12D^; Trp53^fl/fl^; Pd1-Cre* (KP^-/-^C) mice and immortalized pancreatic stellate cells (PSC) cultured in media with or without 1 mM pyruvate (Py) as indicated. The mean value is shown with error bars for standard deviation (n=3). (**b**) Representative NAD(P)H intensity images of PDAC cells and PSCs in monoculture. **(c)** Quantification of optical redox ratio of PDAC cells and PSCs in monoculture (2 independent cultures per cell type were analyzed), normalized to PDAC cells, at single cell level (PDAC cells: n=227; PSCs: n=265). **(d)** For imaging co-cultures, either PDAC cells or PSCs were fluorescently labeled with YFP or GFP, respectively, to distinguish each cell type. Two distinct co-cultures were prepared with one labeled and one unlabeled cell type and optical redox ratios were assessed in the unlabeled cell type to circumvent interference of fluorescent labels with FAD signal due to spectral overlap. For both cultures, PDAC cells were mixed with PSCs in a 1:1 ratio and cultured in standard 2D conditions for 24 hours prior to assessment of the optical redox ratio. Left panel shows representative NAD(P)H intensity (white) images of unlabeled PDAC cells in co-culture with GFP+ PSCs (labeled PSC, green). Right panel shows representative NAD(P)H intensity (white) images of PDAC cells labeled with YFP (labeled PDAC, yellow) in co-culture with unlabeled PSCs. (**e**) Quantification of optical redox ratio of PDAC cells and PSCs from 2 independent cultures per condition as shown in (d). Optical redox ratio was computed for only unlabeled cells, at single cell level (PDAC cells: n=91; PSCs: n=119) and normalized to PDAC cells. Error bars in (c) and (e) are the 95% confidence interval. Statistical significance for (a), (c), and (e) were accessed using one-way ANOVA with Tukey’s posthoc test (*** p< 0.001; ** p< 0.01; * p< 0.05).

To determine whether the redox differences between PDAC cells and PSCs are altered by co-culture, we took advantage of the fact that the optical redox ratio, measured using autofluorescence of NAD(P)H and FAD (i.e., I_NAD(P)H_/I_FAD_), allows assessment of redox state in different cell types in a mixed population. First, we confirmed that the optical redox ratio reflected the differences in NADH/NAD^+^ ratio observed in PDAC cells and PSCs in 2D monoculture (Figure 2b-c). Next, we performed optical redox imaging of PDAC cell-PSC co-cultures. Interestingly, we find that the redox differences observed in PDAC cells or PSCs grown in monoculture were also are present in co-culture such that PDAC cells are more reduced and PSCs are more oxidized; however, the optical redox ratios of the PDAC cells and PSCs in 2D co-culture were closer to each other than the optical redox ratio observed for each cell type in monoculture (Figure 2d-e, Supplementary Figure 3a). A difference in optical redox ratio between PDAC cells and PSCs was also observed when cells were imaged after 48 hours in monoculture, and co-culture still had little to no effect on the redox state of PSCs (Supplementary Figure 3b and 3c). A difference in optical redox ratio was also observed between PSCs and PDAC cells in both mono- and co-cultures when the cancer cells were derived from the widely used KPC autochthonous PDAC mouse model (Supplementary Figure 3d, e). Taken together, these data suggest that PSCs are more oxidized than cancer cells and demonstrate that optical redox imaging can provide insight into the cellular metabolic states of different cell types in a mixed cell population.

### PDAC cells and PSCs exhibit redox differences in 3D organoid co-cultures

We next questioned whether the differences in redox state between PDAC cells and PSCs were retained in 3D organoid co-cultures where PSCs can replace pyruvate addition to the media to support growth. In addition, because optical redox ratio imaging is non-destructive and amenable to analysis of 3D samples at single cell level, we assessed the redox states of different cells in organoid co-cultures over time, with and without pyruvate addition to the culture media. Quantification of organoid number and size shows an increase in both parameters over time in all the culture conditions evaluated (Figure 3a-c, Supplementary Figure 4a, 4b). In organoid cultures, PDAC cells also exhibit a more reduced optical redox ratio compared to PSCs at all time points regardless of whether the different cells types were grown in monoculture, or co-culture, or whether pyruvate was included in the medium (Figure 3d,e; see also Supplementary Table 1). These data argue that cancer cells in organoid cultures are in a more reduced metabolic state than PSCs.

**Figure 3.**
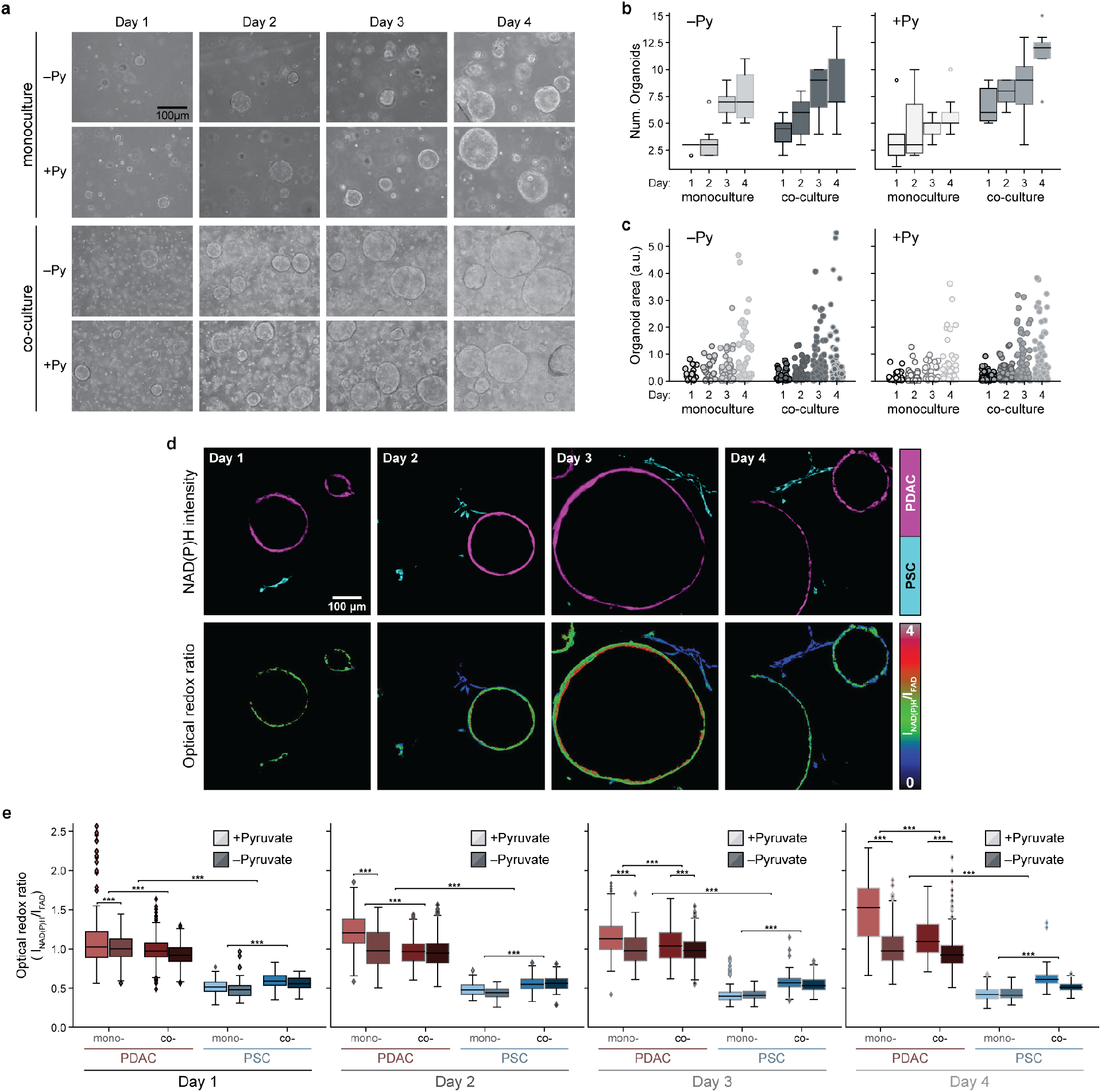
The differences in redox state between PDAC cells and PSCs are diminished in organoid co-culture. **(a)** Representative brightfield images of PDAC cells cultured as 3D organoids in monoculture or in co-culture with immortalized PSCs (PSC), in DMEM supplemented with or without 10mM exogenous pyruvate (Py) as indicated. Images are shown from days 1 through 4 after plating. The number of organoids (Num.Organoids) formed **(b)** and the area for each organoid **(c)** was quantified from brightfield images of 3 independent cultures per condition on each day as indicated. Data shown is the average value per image per condition (n=4 images analyzed per condition). Statistical significance for b and c was accessed using one-way ANOVA with Tukey’s posthoc test and these same data are presented differently in Supplementary Figure 4a-b for all the conditions. **(d)** Representative NAD(P)H fluorescence intensity images of PDAC cells cultured as 3D organoids (shown in magenta) and PSCs (shown in cyan) from co-cultures obtained on days 1 through 4 after plating as indicated (top). The corresponding optical redox ratio from the same images is also shown (bottom). **(e)** Quantification of optical redox ratios of PDAC cells cultured as 3D organoids or PSCs cultured in 3D either in monoculture (mono-) or co-culture (co-) for days 1 through 4. The cultures were grown in DMEM supplemented with or without 10mM exogenous pyruvate (Py). Optical redox ratio is analyzed for n=6 images from the 3 independent cultures per condition per day. The data is computed at single cell level and normalized to PDAC cells in monoculture, grown without exogenous pyruvate for each day. The number of cells segmented per condition is presented in Supplementary Table 1. Statistical significance was accessed using one-way ANOVA with Tukey’s posthoc test (*** p< 0.001; ** p< 0.01; * p< 0.05). All error bars are the 95% confidence interval.

The difference in optical redox ratio between mono- and co-culture conditions with pyruvate show that cancer cells in organoid co-culture with PSCs become more oxidized by day 4 compared to cancer cells in organoid monoculture (Figure 3e). Conversely, PSCs in co-culture conditions with pyruvate become more reduced on day 4 compared to the monoculture condition (Figure 3e). The difference in optical redox ratio between PDAC cells and PSCs is also greater in monoculture than in co-culture for each day (Supplementary Figure 4c), suggesting that PSCs may promote a more oxidized state in cancer cells.

The optical redox differences between PDAC cells and PSCs observed were similar when, instead of imaging the same dish daily for 4 days as shown in Figure 3, the cells were plated as 3D organoid mono- and co-cultures every 24 hours and all the cultures were imaged on day 4 (Supplementary Figure 4e; see also Supplementary Table 2). These data suggest that the measured differences in redox ratio are present across multiple biological replicates. The pyruvate and lactate in the 3D culture media were also assessed on Day 3 and Day 4. Of note, the optical redox ratio of 3D PDAC cells in monoculture correlated with the pyruvate to lactate ratio present in the culture media, while the optical redox ratio of both cell types was decoupled from the pyruvate to lactate ratio in co-culture (Supplementary Figure 4f). Select additional metabolites in the 3D culture media were also measured on day 3 and day 4, but no changes observed were sufficient to explain how PSCs might affect the redox state of cancer cells (Supplementary Figure 5, Supplementary Table 3).

### Physical contact with PSCs facilitates redox changes in cancer cells

PSCs were noted to spread towards the PDAC cells grown as 3D organoid co-cultures, suggesting that PSCs wrap around the PDAC cell organoids or form bridges between different organoids by day 3 and day 4, respectively (Figure 3d). Of note, there is evidence that mechanical interactions between PDAC cells and some populations of PSCs can provide some of the growth promoting benefits of co-culture^4^, although whether cell-cell interactions may be modulating a change in redox state is unknown. Thus, to begin to investigate the possibility that physical contact with PSCs might influence the redox state of PDAC cells, we questioned whether the redox changes could be recapitulated in organoids cultured in media with exogenous pyruvate supplementation or in PSC-conditioned media (derived from PSCs in 3D monoculture). We performed optical redox imaging of PDAC cells grown as organoids in these culture conditions as well as in co-culture with PSCs. Organoids in co-culture with PSCs were greater in number and area than organoids grown with exogenous pyruvate supplementation or in PSC-conditioned media (Figure 4a and b). Compared to the organoids grown with exogenous pyruvate, the organoids grown in conditioned media had a significantly lower optical redox ratio, indicating a more oxidized state, and this ratio in cancer cells was lower still when PSCs were added to the 3D culture system (Figure 4c). This further supports that PSC-derived factors, in addition to pyruvate, may be responsible for cancer cell redox changes. Furthermore, the observation that co-culture with PSCs had a larger impact on the optical redox ratio than conditioned media (Figure 4c), argues that the physical presence of PSCs and cell-cell interactions may contribute to the effects on redox state.

**Figure 4.**
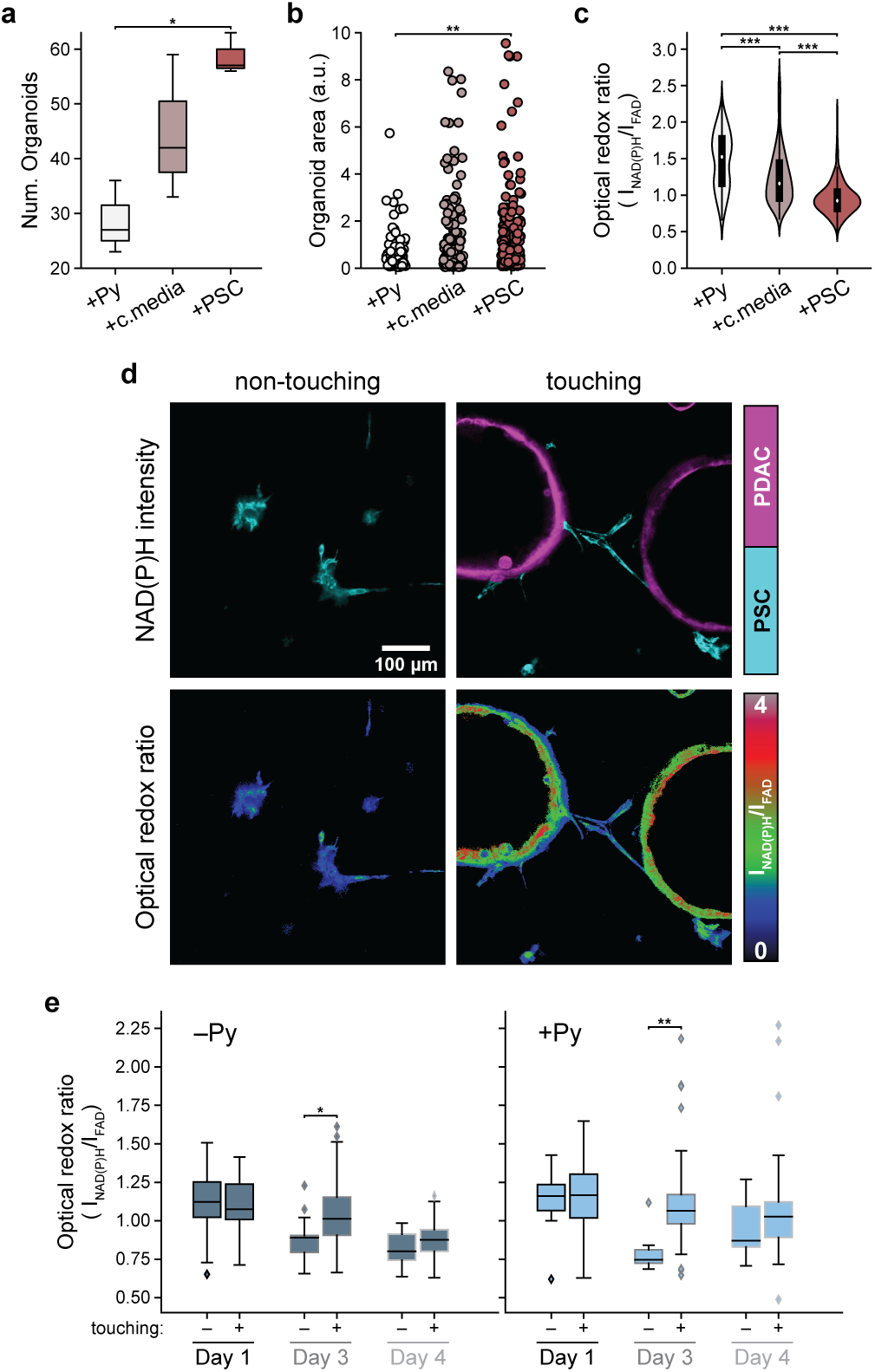
Physical interactions between PSCs and PDAC cells influence the redox state of both cell types and organoid growth. **(a)** PDAC cells cultured as 3D organoids were subject to three different conditions: in media with 10mM exogenous pyruvate (+Py) for 4 days; DMEM media for 2 days followed by addition of PSC conditioned media (+c.media) for 2 days; or in co-culture with PSCs (+PSC) for 4 days. The number of organoids formed **(a)** and the area for each organoid **(b)** was quantified from brightfield images of 3 independent cultures per condition (n=6 images per condition). Each datapoint is one organoid. The differences measured were statistically significant as determined using one-way ANOVA with posthoc Tukey’s test. **(c)** The optical redox ratio for all three conditions described in (a) was computed for each PDAC cell segmented from the organoids and normalized to PDAC cells without pyruvate. Number of cells segmented: n(+Py) = 201; n(+c.media) = 329; n(+PSC) = 347, from n=6 images acquired from the same 3 independent culture dishes as (a-b). Some of this data is a subset of data shown in Figure 3e. Statistical comparisons were done using one-way ANOVA with posthoc Tukey’s test (*** p< 0.001; ** p< 0.01; * p< 0.05). **(d)** Representative NAD(P)H intensity images of PSC cells (shown in cyan) identified as non-touching (left) and touching (right) depending on their proximity to the PDAC cells cultured as organoids (shown in magenta) in co-cultures (top panel). Corresponding optical redox ratio images are also show for each image (bottom panel). PSCs within the 454 μm field-of-view of each organoid were assigned to the touching group, with remaining PSCs assigned to the non-touching group. **(e)** Optical redox ratio computed for PSCs separated into non-touching (indicated by ‘-’) and touching groups (indicated by ‘+’) for day 1, 3, and 4 after plating with and without 10mM exogenous pyruvate (Py). n=6 images were quantified from 3 independent cultures. The data is computed at single cell level and normalized to PDAC cells in monoculture, grown without exogenous pyruvate for each day. The number of cells segmented per condition is presented in Supplementary Table 2. The statistical significance was assessed using T-test (*** p< 0.001; ** p< 0.01; * p< 0.05).

To further investigate a role for cell-cell interactions in modulating redox state, PSCs were divided into ‘touching’ and ‘non-touching’ subsets depending on their spatial proximity to the organoids (Figure 4d and e). The single cell level optical redox ratio was calculated for PSCs in each group for day 1, day 3, and day 4 of co-culture to capture time points that were clearly before and after the observed wrapping of PSCs around organoids. There was no difference in optical redox ratio between the two groups on day 1 (Figure 4e). However, by day 3 the PSCs in the touching group had a significantly more reduced redox state compared to the non-touching group (Figure 4e). On day 4, the redox differences appear to be maintained, although this difference was no longer statistically significant. Overall, these results suggest that co-culture with PSCs supports PDAC cell proliferation through secretion of factors into the media as well as by cell-cell interactions such that the PSCs in contact with cancer cells assume a more reduced redox state than those not in contact with cancer cells. Taken together, these data suggest that PSCs engage in a metabolic interaction with PDAC cells to promote a more oxidized state in the cancer cells.

### Redox differences between different cells are also found in pancreatic tumors

To explore whether the redox differences observed with *in vitro* culture systems are also reflective of the redox state of different cell types in pancreatic tumors, we used the KP^-/-^C and KPC pancreatic cancer mouse model to image redox state in tumor tissue *in vivo* (in live animals) and *ex vivo* (freshly excised tumor tissue). A heterogenous FAD intensity distribution was observed across cell types within tumors imaged both *in vivo* and *ex vivo* (Figure 5a). To identify the number of different cell populations with heterogenous FAD intensity, gaussian fitting was performed on FAD intensity histograms (Figure 5b), which resulted in a three gaussian fit. The first gaussian (Gauss1, gray) was attributed to low intensity pixels from background while the second (Gauss2, blue) and third (Gauss3, red) gaussians indicated the presence of two cell populations. A scatter plot of FAD versus NAD(P)H intensity shows two populations of cells, one with high FAD intensity (FAD_high_) and one with low FAD intensity (FAD_low_) separated at threshold (black dotted line in Figure 5c) calculated from gaussian fits (Figure 5b). The optical redox ratio calculations of bulk tumor tissue (FAD_low_) and interspersed high FAD intensity cells (FAD_high_) shows a more oxidized optical redox ratio for the interspersed cells, with similar observations made when tumor tissue was imaged *in vivo* or *ex vivo* (Figure 5d).

**Figure 5.**
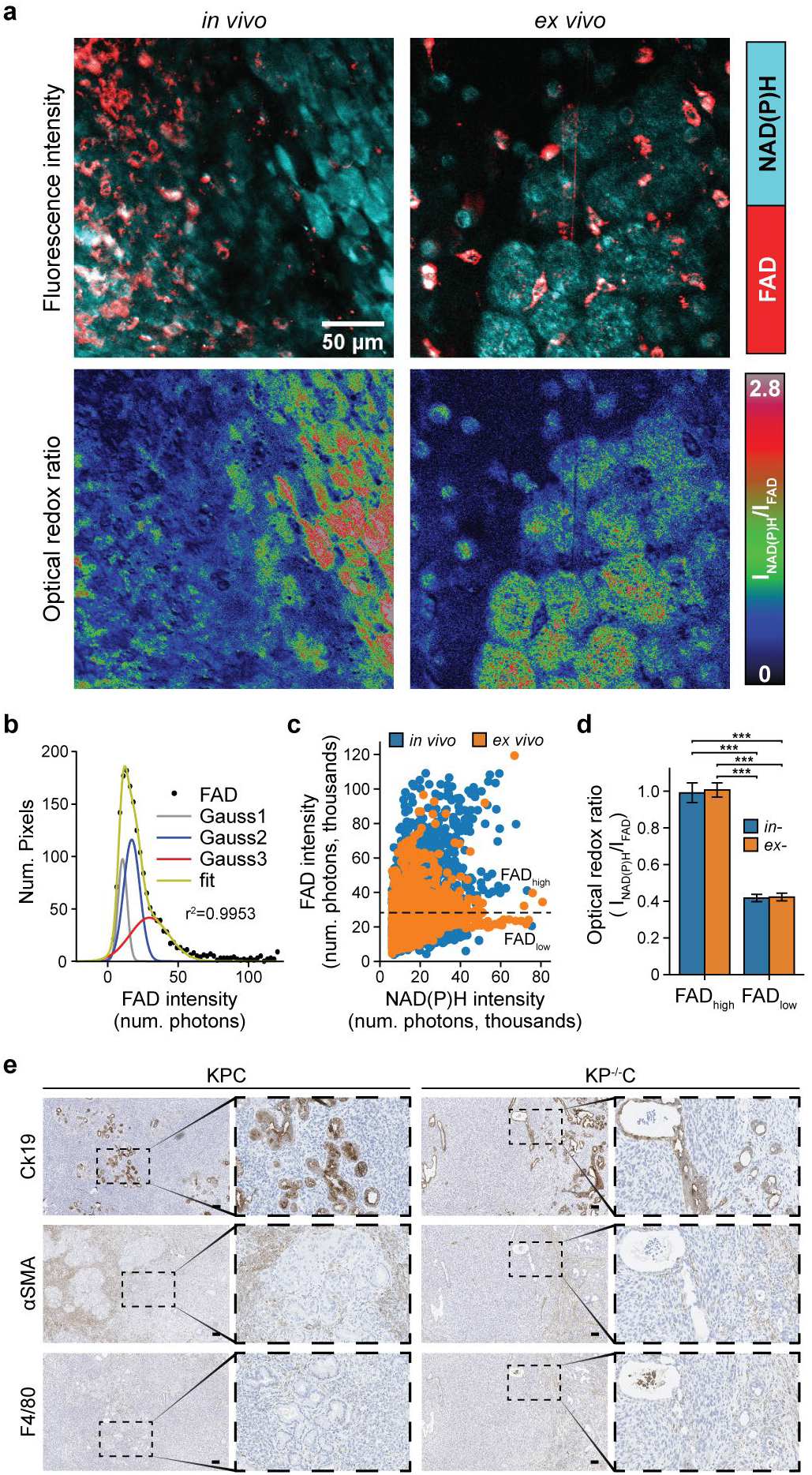
Cells in pancreatic tumors exhibit distinct redox states. (**a**) Representative NAD(P)H (cyan) and FAD (red) intensity images (top), and corresponding optical redox ratio images (bottom), obtained from imaging *in vivo* and freshly excised (*ex vivo*) pancreatic tumor tissue. Imaging was performed on tumors and tumor tissue from *LSL-KRas^G12D^; Trp53^R172H/+^; Pdx1-Cre* (KPC) and *LSL-KRas^G12D^; Trp53^fl/fl^ Pdx1-Cre* (KP^-/-^C) mice. (**b**) Gaussian fit (green curve) was performed on FAD intensity (black solid circles) with the goodness of fit of (r^2^ = 0.9953). Gray (Gauss1), blue (Gauss2) and red (Gauss3) curves represent the best 3-component gaussian fits. The intersecting point between Gauss2 and Gauss3 was computed to be 25.3 × 10^3^ photons. (**c**) Scatter plot of FAD versus NAD(P)H intensity computed from 9 *in vivo* and 8 *ex vivo* tissue images as indicated (n=4 mice). Each datapoint are regions of interest (ROIs) segmented from NAD(P)H and FAD intensity images, n (*in vivo*) =3990, n (*ex vivo*) = 4647. Black dotted line is drawn at 25.3 × 10^3^ photons, the intersecting point calculated in (b) and serves as the threshold between high FAD intensity regions (FAD_high_) and low FAD intensity regions (FAD_low_). (**d**) Optical redox ratio from *in vivo* and *ex vivo* images for regions within FAD_low_ (bulk tissue, low FAD intensity) and FAD_high_ (interspersed high FAD intensity cells) (n=17 images analyzed). The optical redox ratio for each image was normalized to its FAD_high_ regions. Statistical significance was tested using one-way ANOVA with posthoc Tukey’s test. (*** p< 0.001; ** p< 0.01; * p< 0.05). Error bars are the standard deviation. (e) Representative IHC images of PDAC tumors from KPC and KP-/- C mice stained for the PDAC cell marker CK19 (top), fibroblast marker αSMA (middle), or macrophage marker F4/80 (bottom). Scale bars represent 100μm.

Mouse models were not available with a specific label for PSCs in the far red so as not to interfere with the weak signals from NAD(P)H and FAD fluorescence, and fluorescent antibodies against PSCs are difficult to deliver *in vivo* and achieve distribution throughout tumor tissue^60^. Therefore, we processed parallel tumors from the same mouse model to compare how different cell populations are distributed in tumors for comparison to the cell redox distributions observed upon redox imaging of tumor tissue. Histological analysis confirms heterogeneous tumors including cell populations that stain positive for markers of macrophages, pancreatic stellate cells, and epithelial cells (Figure 5e). Of note, stromal cells are distributed among nests of cancer cells. This is consistent with the dispersed FAD high-cell population representing a stromal cell population in the tumors. When considered along with the findings in organoid co-cultures, these data suggest that pancreatic tumors are composed of cell types with distinct redox states and support the hypothesis that stromal cells are more oxidized than cancer cells.

## Discussion

The tumor microenvironment impacts pancreatic tumor development and progression. Pancreatic tumor organoid model systems that incorporate stromal cells can be used to study the cooperative interactions between cancer cells and the supporting stromal cells^2,4,9^. Here we take advantage of these systems and optical redox imaging and find that PSCs assume a more oxidized redox state than cancer cells and may engage in redox interactions with cancer cells to support proliferation. This interaction could help pancreatic cancer cells carry out oxidation reactions despite limited access to oxygen as an electron acceptor, and direct contact between cancer cells and PSCs appears necessary to maximize this effect.

Optical redox imaging has been previously used to study metabolic interactions between HeLa cells and human skin fibroblasts^61^. Consistent with our findings, the fibroblasts were more oxidized compared to cancer cells in 2D monoculture. Optical redox imaging also revealed a more oxidized state for the human MIA PaCa-2 PDAC cell line when co-cultured in 3D with patient-derived CAFs compared to monoculture conditions, but did not assess the redox state of the CAFs^62^. Nevertheless, these studies further support a hypothesis where interactions between cancer cells and stromal cells can promote a more oxidized state in cancer cells.

Access to electron acceptors can limit cancer cell proliferation, as NAD+ regeneration supports oxidized biomass production^23–25,27,28,31,63^. Consistent with this notion, we find cancer cells in both organoid and mouse tumor PDAC models are more reduced than surrounding non-cancer cells. Gaining access to exogenous electron acceptors, such as pyruvate, can overcome this redox limitation^23,24^, and secretion of pyruvate by PSCs is one mechanism by which stromal cells might help cancer cells proliferate^58^. However, the concentration of pyruvate in interstitial fluid isolated from pancreatic tumors is lower than the concentration of pyruvate found in plasma, suggesting that some cells in pancreatic tumors may not have extensive access to this metabolite *in vivo*^64^. Nevertheless, local exchange of pyruvate and lactate between cells is possible and would not be evident in bulk fluid measurements. It is also conceivable that NADH/NAD^+^ ratio varies among different cancer cell populations within tumors and over different stages of disease progression, in alignment with nutrient fluctuations within the tumor microenvironment or based on interactions between cancer cells and stromal cells that vary in both time in space.

In addition to secretion of pyruvate or other metabolites, our findings argue that physical interactions between the cancer cells and the stellate cells also help cancer cells overcome redox limitations of proliferation. As previously reported, physical interactions between pancreatic stellate cells and PDAC organoids can support proliferation^4^ and induce a pro-inflammatory state in the stromal cells. How physical interaction with PSCs might alter the redox state of cancer cells as well as how PSCs maintain a more oxidized state than cancer cells is unknown. Close contact may enable local exchange of pyruvate and lactate or another pair of metabolites to transfer electrons between cell types. The coupling of hydrophobic molecules in plasma cell membranes to transfer electrons is another possibility for how cell contact might mediate redox changes in different cell types, and there is evidence of electron transport in cell membranes other than the mitochondrial membrane, including the plasma membrane^65^. In fact, plasma membrane electron transfer has been reported in pancreatic cells^66^, and may be involved in how physical interaction with PSCs maintains a more oxidized state in pancreatic cancer cells.

Mechanical forces including those produced by cell-cell interactions and interactions with a more or less stiff extracellular matrix (ECM) can also affect cell metabolism, and fibroblasts are known for their ability to synthesize the protein that comprise the ECM component of the pancreatic tumor stroma ^67,68^. The ECM can influence nutrient uptake from the environment as well as affect cell signaling pathways that regulate metabolism^67^. Metabolite exchange has also been reported between cell types in pancreatic cancer cells based on physical contact. For example, cell-cell interactions between the glutamatergic pre-synaptic protein Netrin G1 in CAFs and its receptor NGL-1 in PDAC cells can play a role glutamate/glutamine sharing between cell types^69^. Regardless of the mechanism, when taken together the data presented argue that direct interactions between PSCs and cancer cells can promote a more oxidized state in the cancer cells, and suggest that redox sharing is another way by which cell-cell interactions between cancer cells and stromal cells can support pancreatic cancer metabolism and tumor growth.

## Materials and methods

### Cell culture

To generate PDAC cell lines or organoids for *in vitro* experiments, cancer cells were isolated from pancreatic tumors arising in the *Kras^+/LSL-G12D^; Trp53^+/LSL-R172H^; Pdx1-Cre (KPC)* or *Kras^+/LSL-G12D^; Trp53^-/-^; Pdx1-Cre (KP^-/-^C)* mouse models bred onto a C57BL/6J background using established protocols^9^. In brief, PDAC tumors were harvested, minced, and digested in 5 mL of PBS containing cold 1 mg/mL collagenase I (Worthington Biochemical LS004194), 3mg/mL dispase II (Roche 04942078001) and 0.1 mg/mL DNase I (Sigma D4527) rotating at 37° C for 30 min. After digestion, EDTA was added to a final concentration of 10 mM and the solution was filtered through a 70 μm cell strainer and then incubated at room temperature for 5 min before spinning at 200 RCF for 5 min at 4° C and washed with PBS. For cell lines, cells were resuspended in DMEM containing pyruvate with 10% heat inactivated serum and penicillin-streptomycin.

To isolate and immortalize PSCs, GFP^+^ PSCs were isolated from *β-actin-GFP* mice on a C57Bl6/J background (006567) as previously described^9,70,71^. Briefly, 3 mL of 1.3 mg/mL cold collagenase P (Sigma 11213865001) and 0.01 mg/mL DNase (Sigma D5025) in GBSS (Sigma G9779) were injected into the pancreas. The tissue was then placed into 2 mL of collagenase P solution on ice. Cells were then placed in a 37° C water bath for 15 minutes. The digested pancreas was filtered through a 250 μm strainer and washed with GBSS with 0.3% BSA. A gradient was created by resuspending the cells in Nycodenz (VWR 100356-726) and layering in GBSS with 0.3% BSA. Cells were then centrifuged at 1300g for 20 minutes at 4° C. The layer containing PSCs was removed, filtered through a 70 μm strainer, washed in GBSS with 0.3% BSA, and plated for cell culture in DMEM with 10% FBS and penicillin-streptomycin. GFP^+^ PSCs were immortalized with TERT and SV40 LargeT after several passages. Unlabeled, immortalized PSCs were a gift from Richard Hynes’s and David Tuveson’s laboratories^4^. Immortalized mouse embryonic fibroblasts (MEFs) were created as described previously^72^. Cell lines were regularly tested for mycoplasma contamination using the MycoAlert Plus kit (Lonza) or the Mycoprobe Mycoplasma Detection Kit (R&D Systems).

### Organoid culture

To generate organoids for *in vitro* experiments, cells were isolated from pancreatic tumors arising in *Kras^+/LSL-G12D^; Trp53^-/-^; Pdx1-Cre* (*KP^-/-^C*) mouse models bred onto the C57BL/6J background using established protocols^9,20^. Cancer cells were isolated directly from mice bearing PDAC tumors and cultured as organoids, referred to here as PDAC cells cultured as 3D organoids. Organoids were never exposed to standard 2D culture conditions and were propagated as organoids as described previously^9,20^. PDAC cells were resuspended in 50 μL growth factor reduced (GFR) Matrigel (Corning 356231) and propagated in organoid feeding media. Specifically, organoid domes were cultured in 500 μL complete feeding media that consisted of Advanced DMEM/F12 (Thermo Fisher 12634) containing GlutaMAX (Thermo Fisher 35050), penicillin-streptomycin, HEPES (Thermo Fisher 15630), 0.5 μM TGF-b inhibitor A-83-01 (TOCRIS 2939), 0.05 μg/mL EGF (Thermo Fisher PMG8041), 0.1 μg/mL FGF (Peprotech 100-26), 0.01 μM Gastrin I (TOCRIS 3006), 0.1 μg/mL Noggin (Peprotech 250-38), 10.5 μM Rho Kinase Inhibitor Y-27632 (Sigma Y0503), 1.25 mM N-Acetylcysteine (NAC) (Sigma A9165), 10 mM Nicotinamide (Sigma N0636), 1X B-27 supplement (Thermo Fisher 17504), and 1 μg/mL R-spondin. R-spondin was purified from 293T cells engineered to produce it using a Protein A Antibody Purification Kit (Sigma PURE1A). For PDAC cells grown as 3D organoids in co-culture, co-cultures were grown in DMEM without pyruvate (Corning 10-017-CV) supplemented with 10% dialyzed FBS and penicillin-streptomycin (minimal media). PDAC cells grown as 3D organoids were regularly tested for mycoplasma contamination using the MycoAlert Plus kit (Lonza) or the Mycoprobe Mycoplasma Detection Kit (R&D Systems).

For passaging and assessment of cell numbers, PDAC cells grown as 3D organoids were digested to single cells by incubating with 2 mg/mL dispase in Advanced DMEM/F12 with penicillin-streptomycin, HEPES, and GlutaMAX at 37° C for 20 minutes. PDAC cells grown as 3D organoids were then triturated with a fire-polished glass pipette and enzymatically digested with 1 mL TrypLE Express (Thermo Fisher 12605-010) for 10 minutes rotating at 37° C, followed by addition of 1 mL of dispase containing media and 10 μL of 10 mg/mL DNAse (Sigma 4527) and digested rotating at 37° C for 20 minutes or until single cells were visible under a microscope. PDAC cells were counted and plated in GFR Matrigel at a concentration of 2000 cells/well.

For co-culture experiments using transwells, 2000 PDAC cells grown as 3D organoids were plated on the bottom of the plate in a Matrigel dome and 10,000 pancreatic cancer cells, MEFs, or PSCs were plated on the top of the transwell insert in 24 well plates. Organoids were imaged on day 6 after plating.

For co-culture experiments of PDAC cells cultured as 3D organoids and stromal cells including addition of exogenous pyruvate or alanine, 2000 PDAC cells and 10,000 MEFs or PSCs were mixed together and plated in a Matrigel dome in 24-well plates. 1 mM alanine or 1 mM or 10 mM pyruvate was added to the culture media, and cells were imaged on day 6 after plating.

### Sample preparation for multiphoton imaging

For multiphoton imaging of 2D monocultures, either PDAC cells isolated from the KP^-/-^C or KPC mouse models or PSCs were seeded (seeding density: 200,000) on 35 mm glass-bottom dishes (#P35G-1.5-14-C, MatTek) with 2mL media (DMEM with 10% FBS and 1% pen/strep), either 24 hours or 48 hours before imaging. For 2D co-cultures, either fluorescently labeled PDAC cells from the KP^-/-^C model (labeled with YFP), PDAC cells from the KPC model (labeled with tdTomato) or PSCs (labeled with GFP) were used to distinguish each cell type during imaging. Of note, the optical redox ratio of cells labeled with GFP, YFP and tdTomato cannot be measured since these fluorophores obscure the FAD autofluorescence signal due to spectral overlap. Thus, data was obtained from distinct sets of co-cultures in this study where only one of the two cells types was labeled. For all co-cultures, the two cell types were mixed in 1:1 ratio and cultured for either 24 hours or 48 hours before imaging.

Multiphoton imaging of 3D culture of organoids and PSCs was performed in 35 mm glass-bottom dishes. PDAC cells cultured as organoids or PSCs were embedded in a Matrigel matrix and overlaid with media (DMEM without pyruvate supplemented with 10% dialyzed FBS and 5% penicillin-streptomycin). For co-culture, PSCs (seeding density: 200,000 cells/dish) were mixed with the PDAC cells. For +pyruvate conditions, 10mM pyruvate (Sigma) was added to the media. At least three representative organoids were imaged near the center of the organoid per dish.

### Bright field imaging and analysis

All bright field images were collected on a Nikon Eclipse Ti-S microscope using 2X, 0.10 N.A or 4X, 0.13 N.A objectives, except data for Figure 3a-c and Figure 4a-b were collected on an EVOS XL CORE Imaging microscope (ThermoFisher Scientific) using 20X, 0.40N.A (LPlan PH2, EVOS) and 4X, 0.13 N.A (LPlan PH2, EVOS) objectives respectively. Image analysis, organoid segmentation, organoid area and number calculations were performed using ImageJ software (NIH)^73^.

### Multiphoton Imaging and optical redox ratio analysis

#### 2D and 3D cultures

Multiphoton imaging of 2D and 3D cultures was performed on a custom-built Ultima Multiphoton Imaging System (Bruker) consisting of an inverted microscope (TI-E, Nikon) coupled to an ultrafast tunable laser source (Insight DS+, Spectra Physics). The data was acquired in photon counting mode using time correlated single photon counting electronics (SPC 150, Becker & Hickl GmbH). Imaging was performed using Prairie View Software (Bruker). The tunable multiphoton laser allowed sequential excitation of NAD(P)H and FAD/YFP/GFP/tdTomato at 750 nm and 890 nm, respectively. All 2D culture samples were illuminated through a 40x water immersion, 1.15 N.A objective (Plan Apo, Nikon) with an image scan speed of 4.8 μs/pixel, 60 second integration time, image size of 256 × 256 pixels, and field of view of 270μm. 3D culture samples were imaged via a 20X air objective, 0.75 N.A (Plan Apo, Nikon) with 1.19X digital zoom, image scan speed of 2.4 μs/pixel, 100 second integration time, image size of 512 × 512 pixels, and field of view of 454 μm. A dichroic at 720 nm was used to separate excitation from emission signals. The emission filter used for NAD(P)H was bandpass 460/80 nm, while FAD, YFP and GFP were collected using bandpass 550/100nm. GaAsP photomultiplier tubes (H7422P-40, Hamamatsu, Japan) were used for detection.

For the data collected on photon counting mode (2D and 3D cultures), the fluorescence decay curve at each pixel was integrated to generate the intensity of each fluorophore using SPCImage software (Becker & Hickl GmbH). For all data, fluorescence intensity image analysis was performed using ImageJ software (NIH)^73^. For single cell analysis, a customized CellProfiler pipeline was used to segment individual cell cytoplasms^74^. Cytoplasm masks were applied to all images to determine single-cell optical redox ratio using equation (1) where *I* represents the intensity of the fluorophore at each pixel.

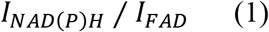

For 2D co-cultures, the optical redox ratio from only unlabeled cells in co-cultures were computed by eliminating cells with GFP, YFP or tdTomato fluorescence signal in the imaged region.

#### In vivo and ex vivo samples

All animal work was done in accordance with the guidelines and procedures approved by Massachusetts Institute of Technology Committee on Animal Care. For *in vivo* imaging, the mice were anaesthetized by intraperitoneal injection of Avertin. A small vertical incision on the left side at the level of the pancreas was made followed by pancreatic tumor exteriorization. The exposed organ was orientated so that the tissue was underneath the animal, pressed gently on a glass slide dish for imaging. For *ex vivo* imaging, pancreatic tumor tissue was excised and placed on the glass cover slip of 35 mm glass-bottom dishes. All *ex vivo* imaging was performed within 1 hour after excision.

*In vivo* and *ex vivo* PDAC tumors from mice were imaged on an Olympus multiphoton laser scanning microscope (FV1000MPE) with non-descanned detectors, coupled to an ultrafast Ti:Sapphire laser (DeepSea, Spectral Physics) using 25X water immersion, 1.05N.A objective (Olympus) in analog scanning mode. Image acquisition speed was 12.5μs/pixel, and total integration time was 118 seconds. The image size was 512 × 512 pixels with a field of view of 254μm. The emission filters used for NAD(P)H and FAD were bandpass 472/30 nm and 525/45nm, respectively.

Image analysis for *ex vivo* and *in vivo* experiments was performed on ImageJ software. Regions of interest (ROI) were segmented from NAD(P)H intensity images and masks were generated using ‘analyze particles’ function. Similarly masks for FAD_high_ ROI were generated from thresholded FAD intensity images. FAD_low_ ROI masks were generated by applying a XOR function between original NAD(P)H ROI and the FAD_high_ ROI masks. To generate consistent number of ROIs per image, the images were subsampled into 1024 images of size 16 by 16 pixels. The optical redox ratio for each segmented ROI was calculated using equation (1).

### Biochemical assessment of NADH/ NAD^+^ ratio

NAD^+^/NADH measurements were done using a modified NADH/ NAD^+^ Glo Assay (Promega) as described previously (Sullivan et. al., 2015). Briefly, 200,000 cells were plated on 35 mm dish in 2 mL media in DMEM. 24 h later, cells were washed 3 times in PBS and cells were maintained in 2 mL of DMEM with or without pyruvate in dialyzed serum for 48 h. For extraction, cells were washed 3 times in ice cold PBS and extracted in 100 μL of lysis buffer (1% Dodecyltrimethylammonium bromide (DTAB) in 0.2 N NaOH diluted 1:1 with PBS) and frozen in −80° C. After establishing the linear range of detection with sample volume and measurement time, NADH was measured by incubating samples at 75° C for 30 min using 5 μL of the lysate and 15 μL of the lysis buffer. An equal sample volume of 5 μL of lysate and 15 μL of lysis buffer was added to 20 μL lysis buffer and 20 μL of 0.4 N HCL and incubated at 60° C for 15 min to measure NAD^+^. Samples were quenched and neutralized with 20 μL of 0.25 M Tris in 0.2 N HCL (NADH) or 20 μL 0.5 M Tris base (NAD^+^) and bioluminescence was measured using a plate reader.

### GC-MS analysis of metabolites

Metabolites were analyzed by gas chromatography-mass spectrometry (GC-MS) as described previously^9,75^. Briefly, 10 μL of each media sample was extracted with 600 μL cold methanol, vortexed at 4° C for 10 min, and spun down at max speed on a table top centrifuge for 10 min at 4° C. 450 μL of each sample was then dried under nitrogen gas and stored at −80° C until GC-MS analysis. Dried metabolite extracts were dissolved in 16 μL methoxamine (MOX) reagent (ThermoFisher TS-45950) and incubated at 37° C for 90 minutes followed by addition of 20μL N–methyl–N–(tert–butyldimethylsilyl)trifluoroacetamide + 1% tert–Butyldimethylchlorosilane (Sigma 375934) and incubated at 60°C for 1 hour. Following derivatization, samples were analyzed using a DB-35MS column (Agilent Technologies) in an Agilent 7890 gas chromatograph coupled to an Agilent 5975C mass spectrometer. Helium was used as the carrier gas at a flow rate of 1.2 mL/min. One microliter of sample was injected at 270°C. After injection, the GC oven was held at 100°C for 1 min. and increased to 300°C at 3.5 °C/min. The oven was then ramped to 320°C at 20 °C/min. and held for 5 min. at this 320°C. The MS system operated under electron impact ionization at 70 eV and the MS source and quadrupole were held at 230°C and 150°C, respectively. The detector was used in scanning mode, and the scanned ion range was 100-650 m/z.

To generate metabolite heatmaps (Supplementary Figure 5), the total ion count (TIC) values of each metabolite was first normalized to Norvaline TIC followed by normalization of each metabolite to that of PDAC cells cultured as 3D organoids without pyruvate. The heatmap score for each metabolite per culture condition was calculated using equation (2)

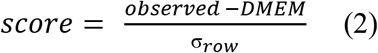

where *observed* is the normalized TIC of the metabolite present in conditioned media from 3D culture, *DMEM* is the TIC of the metabolite present in fresh culture media (DMEM without pyruvate (Corning 10-017-CV) supplemented with 10% dialyzed FBS and 5% penicillin-streptomycin) and *σ_row_* is the standard deviation of the metabolite level of all culture conditions (i.e., heatmap row for the metabolite).

### Immunohistochemistry

5 μM sections from formalin fixed, paraffin embedded mouse tumors were stained with antibodies against cytokeratin-19 (Abcam #ab133496; 1:500 dilution), a-SMA (Cell Signaling Technologies #19245; 1:125 dilution), or F4/80 (Cell Signaling Technologies #70076; 1:125 dilution) using standard techniques.

### Statistical analysis

Data analysis was performed in Python and graphs and heat maps for all the figures were plotted in Python using the Seaborn data visualization library. Statistical tests were performed in R. Data was compared using either Student’s T-test or one-way ANOVA with a Dunnett’s or Tukey’s posthoc test for multiple comparison and p-values have been indicated as follows: *** p< 0.001; ** p< 0.01; * p< 0.05. Gaussian curve fitting for Figure 5b was performed using MATLAB.

## Acknowledgements

We would like to thank the Koch Institute Swanson Biotechnology Center for assistance with microscopy and histology, as well as the members of the Vander Heiden lab and Skala lab for helpful discussions. We would also like to acknowledge Dr. Jens Eickhoff for his advice on statistical analysis, Dr. Steve Trier for guidance with *in vivo* imaging, and Matthew Stefely for input on design of scientific diagrams and data presentation. A.N.L. was a Robert Black Fellow of the Damon Runyon Cancer Research Foundation (DRG-2241-15) and is supported by a NIH Pathway to Independence Award (K99CA234221). S.S. is supported by the Damon Runyon Cancer Research Foundation (DRG-2367-19). M.G.V.H. and M.C.S. received substantial support for this work from SU2C, a division of the Entertainment Industry Foundation. M.G.V.H. acknowledges additional support from the Lustgarten Foundation, a Faculty Scholar grant from the Howard Hughes Medical Institute, the MIT Center for Precision Cancer Medicine, the Ludwig Center at MIT, the Emerald Foundation, and the NCI (R35CA242379, R01CA201276, P30CA14051). M.C.S acknowledges additional support from the NCI (R21CA224280, R01CA211082, R01CA205101, R01CA185747), the NSF (CBET-1642287), the Morgridge Institute for Research, and the University of Wisconsin Carbone Cancer Center.

## Author Contributions

Rupsa Datta: Conceptualization, Methodology, Software, Formal Analysis, Investigation, Writing - original draft, Writing - review and editing, Visualization. Allison N. Lau: Conceptualization, Methodology, Investigation, Writing - original draft, Writing - review and editing. Sharanya Sivanand: Conceptualization, Methodology, Investigation, Writing - original draft, Writing - review and editing. Logan Florek: Formal Analysis. Jeff Wyckoff: Investigation, Methodology. Melissa C. Skala: Conceptualization, Supervision, Funding acquisition, Methodology, Writing - original draft, Writing - review and editing. Matthew G. Vander Heiden: Conceptualization, Supervision, Funding acquisition, Methodology, Writing - original draft, Writing - review and editing.

## Competing Interests

M.G.V.H. is a scientific advisor for Agios Pharmaceuticals, Aeglea Biotherapeutics, iTeos Therapeutics, Faeth Therapeutics, and Auron Therapeutics.

