## Supplementary material for "Interactions with stromal cells promote a more oxidized cancer cell redox state in pancreatic tumors"

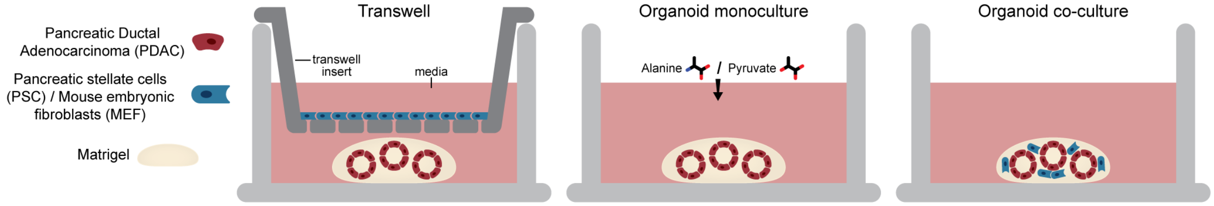


**Supplementary Figure 1**. Schematic representation of transwell, organoid monoculture, and organoid co-culture systems. In the transwell system, PDAC cells cultured as 3D organoids embedded in Matrigel are in the bottom chamber while cells grown in 2D culture are in the upper chamber (transwell insert). In organoid monoculture, PDAC cells alone are cultured as 3D organoids embedded in Matrigel. Where indicated, exogenous alanine or pyruvate is added to the culture medium. In organoid co-culture, PDAC cells are cultured as 3D organoids and mixed with either MEFs or PSCs and embedded in Matrigel.

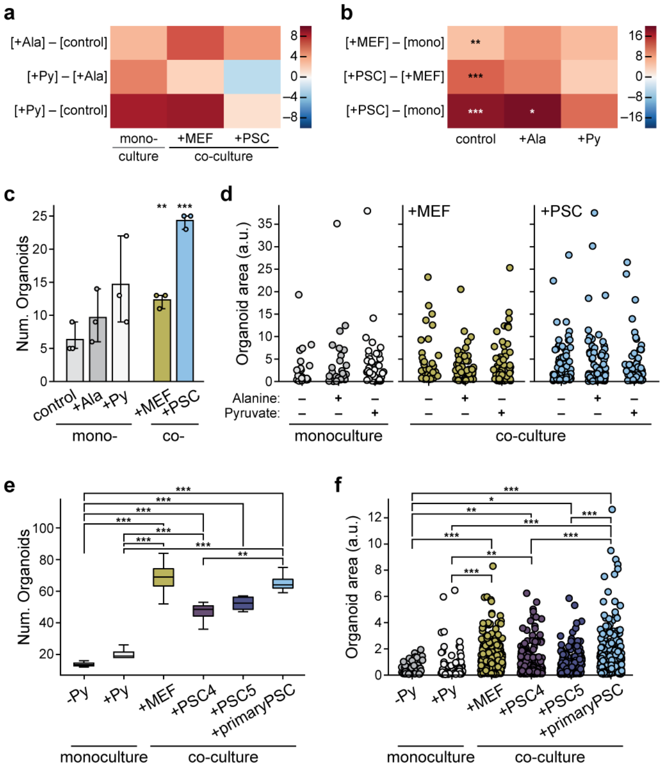

**Supplementary Figure 2**. **(a)** Heatmap of the difference in mean number of organoids for murine PDAC cells cultured as 3D organoids in monoculture, co-culture with MEFs (+MEF), or co-culture with PSCs (+PSC), without (control) or with exogenous 1mM alanine (+Ala) or 1mM pyruvate (+Py) supplemented in the media. First row, [+Ala] – [control], represents the difference in mean organoid number between cultures when media contains 1mM exogenous alanine and media without alanine or pyruvate (control). Second row, [+Py] – [+Ala], represents the difference in mean organoid number between cultures with 1mM pyruvate and 1mM alanine. Third row [+Py] – [control] represents the difference in mean organoid number between cultures with 1mM pyruvate and control. n=3 images were analyzed per condition. Corresponding data are also shown in Figure 1c-d. **(b)** Heatmap of the difference in mean number of organoids derived from murine PDAC cells in mono- and co-cultures grown without any exogenous metabolite (control), or with 1mM exogenous alanine (+Ala) or 1mM pyruvate (+Py). First row, [+MEF] – [mono], represents difference in organoid number between PDAC cells cultured as 3D organoids in co-culture with MEFs and monoculture. Second row [+PSC] – [+MEF], represents the difference in organoid number between co-culture with PSC and co-culture with MEF. Third row, [+PSC] – [mono], represents the difference in organoid number between co-culture with PSCs and monoculture. n=3 images were analyzed per condition. Corresponding data is also shown in Figure 1c-d. **(c)** Number of organoids (Num. Organoids) quantified from brightfield images of PDAC cells cultured as 3D organoid monoculture (mono-) grown in media without exogenous alanine or pyruvate (control), media supplemented with 1 mM alanine (+Ala) or 1 mM pyruvate (+Py). Also plotted are the number of PDAC cells cultured as 3D organoids, co-cultured (co-) with MEFs (+MEF) or PSCs (+PSC) as indicated. 3 independent wells were assessed per condition and each point is the average organoid number measured per well. n=3 per condition. Error bars represent the standard deviation. Data presented here are a subset of the data shown in Figure 1c. **(d)** The area of PDAC cells cultured as 3D organoids alone (monoculture), co-cultured with MEF cells (+MEF) or PSCs (+PSC) with media supplemented with and without 1mM pyruvate and 1mM alanine as indicated. Area of each organoid was quantified from brightfield images (n=3 images per condition). 3 independent wells were assessed per condition. Each data point is an organoid area. **(e)** The number of organoids (Num.Organoids) or **(f)** organoid area of PDAC cells cultured as 3D organoids cultured alone without pyruvate (-Py), alone with 10mM exogenous pyruvate (+Py), co-cultured with MEFs (+MEF), co-cultured with immortalized PSCs (+PSC4, +PCS5), or co-cultured with primary PSCs (+primary PSC). The data is quantified from brightfield images of 4 dishes per condition (n=4 images analyzed per condition). Statistical significance of differences between multiple conditions for data presented in (a-g) were tested using one-way ANOVA with posthoc Tukey's test (*** $p< 0.001$; ** $p< 0.01$; * $p< 0.05$). Error bars for (e) represent the 95% confidence interval.

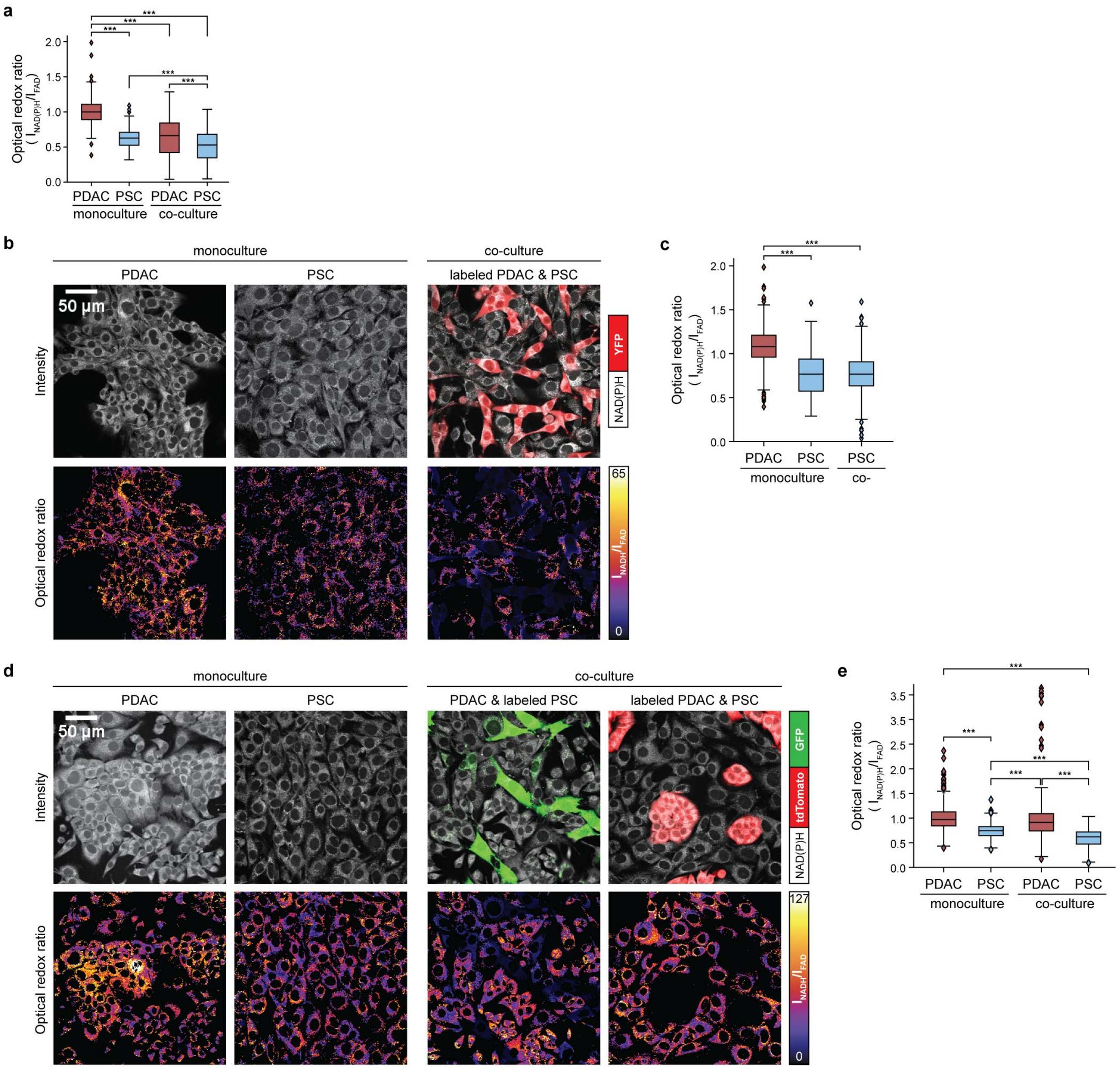


**Supplementary Figure 3**. (a) Optical redox ratio of 2D monocultures and co-cultures of unlabeled murine PDAC cells (KP$^{-/-}$C mouse model) and immortalized unlabeled PSC cells measured 24 hours after plating (2 independent cultures per condition were assessed). The optical redox ratio for all the conditions were calculated at single cell level (PDAC cells monoculture: n=227; PDAC cells co-culture: n=91; PSC cells monoculture: n=265; PSC cells co-culture: n=119). The quantified data was normalized to the optical redox ratio for PDAC cells in monoculture. Data shown are derived from data that is presented in Figure 2 b-e. **(b)** Top panel shows representative fluorescence intensity images of NAD(P)H fluorescence (white) of 2D monocultures of PDAC cells (KP$^{-/-}$C mouse model) and immortalized PSCs (PSC) and 2D co-culture of YFP+ murine PDAC cells (KP$^{-/-}$C mouse model, labeled PDAC) and unlabeled PSCs. Cells were imaged after co-culture for 48 hours, and YFP (red) allowed specification of the cancer cells in co-culture. Corresponding optical redox ratio images are also shown (bottom). **(c)** Quantification of the optical redox ratio of PDAC cells (n=436) and PSCs (n=293) in monoculture, and unlabeled PSCs in co-culture (co-) (n=380) as described in (b) from 2 independent cultures per conditions. The quantified data was normalized to the optical redox ratio for PDAC cells in monoculture. **(d)** Representative fluorescence intensity images of NAD(P)H fluorescence (white) of 2D monoculture of unlabeled PDAC cells (KPC mouse model), monoculture of unlabeled PSCs, co-culture of PDAC cells and GFP+ PSCs (labeled PSC), and co-culture of tdTomato+ PDAC cells (labeled PDAC, KPC mouse model) and PSCs (top), cultured for 48 hours. Two separate co-cultures were prepared with one labeled and one unlabeled cell type, and optical redox ratios were assessed in the unlabeled cell type to circumvent interference of fluorescent labels with FAD signal. GFP intensity is overlaid in green and tdTomato in red with NAD(P)H intensity image in white. The optical redox ratio is also shown for all images (bottom). **(e)** Quantification of optical redox ratio from panel (c). n=8 images for monocultures and n=18-19 images for co-cultures were acquired from 2 independent cultures per condition and normalized to the mean optical redox ratio measured for PDAC cells in monoculture. For co-cultures, the optical redox ratios of unlabeled cells were computed, i.e, unlabeled PDAC cells with labeled PSCs excluded, or unlabeled PSCs with labeled PDAC cells excluded. The optical redox ratio was obtained at a single cell level (PDAC cell monoculture: n=1194; PDAC cell co-culture: n=1307; PSC monoculture: n=642; PSC co-culture: n=874). The statistical significance of differences between conditions shown in (a), (c) and (e) were evaluated using ANOVA with posthoc Tukey's test (*** $p< 0.001$; ** $p< 0.01$; * $p< 0.05$).

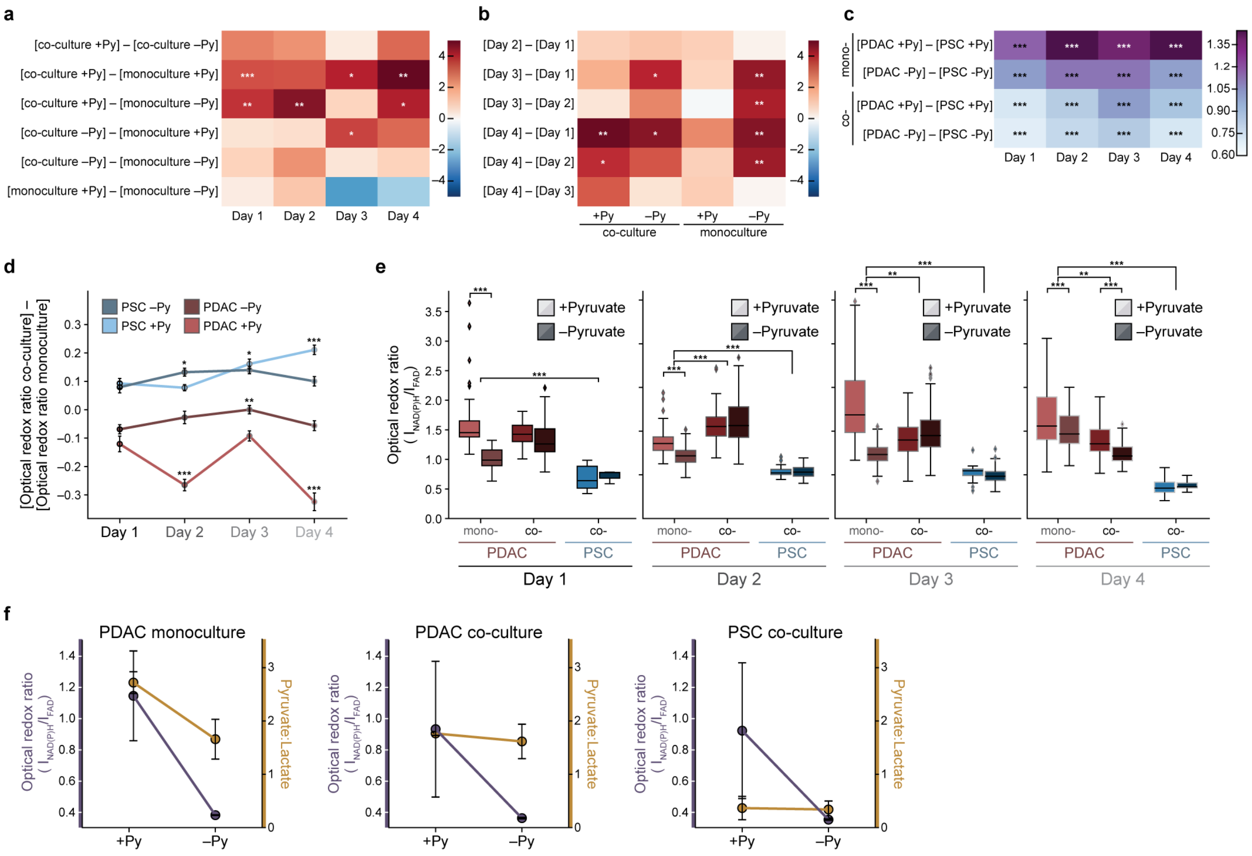


**Supplementary Figure 4. (a)** Heatmap of the difference in means in number of organoids for PDAC cells cultured as organoids from day 1 through 4. Each row represents difference of organoid number between two culture conditions (i.e., either monoculture or co-culture with PSC, supplemented with and without 10mM pyruvate (Py) as indicated). n=4 images were analyzed per condition. Corresponding data are also shown in Figure 3b. **(b)** Heatmap of the difference in mean organoid number for co-culture and monoculture, supplemented with and without 10mM pyruvate (Py) as indicated. Each row represents difference in organoid number between two culture days. n=4 images were analyzed per condition. Corresponding data are also shown in Figure 3b. The statistical significance for (a) and (b) was tested using one-way ANOVA with posthoc Tukey's test (*** $p< 0.001$; ** $p< 0.01$; * $p< 0.05$). **(c)** Heatmap of the difference in means of optical redox ratio between PDAC cells and PSCs [PDAC – PSC], supplemented with or without 10mM pyruvate (Py) and grown as monoculture (mono-) or co-culture (co-) 3D organoids. n=6 images were analyzed per condition. Corresponding data are also shown in Figure 3e. **(d)** Optical redox ratio differences between monoculture and co-culture of PDAC cells cultured as 3D organoids, either supplemented with 10mM exogenous pyruvate or without pyruvate (Py) as indicated. Also plotted are differences in optical redox ratio between monoculture and co-culture of PSCs grown in 3D culture, either supplemented with 10mM exogenous pyruvate or without pyruvate. n=6 images were analyzed per condition. The optical redox ratio differences are plotted for day 1 through 4. The error bars represent the standard error. The data for days 2, 3, and 4 have been compared to day 1 using T-test for each curve (*** $p< 0.001$; ** $p< 0.01$; * $p< 0.05$). The data is derived from the data shown in Figure 3e **(e)** Quantification of optical redox ratios that were computed for each cell segmented from the PDAC cells cultured as 3D organoids in monoculture or co-culture with PSCs from day 1 through 4. Optical redox ratio was measured for the PDAC cells cultured in monoculture (mono-) or co-culture (co-) with the PSCs and PSCs grown in co-culture with the PDAC cells as 3D organoids. 2 dishes per culture condition were plated on each day from day 0 through day 3. All the n= 48 dishes were imaged on day 4 with at least n = 3 images per condition. Optical redox ratios for all the conditions have been normalized to PDAC cells in monoculture on day 1, grown without exogenous pyruvate. The statistical significance was tested using one-way ANOVA with posthoc Tukey's test (*** $p< 0.001$; ** $p< 0.01$). **(f)** Optical redox ratio (gray) and pyruvate to lactate ratio measured in media (yellow) for PDAC cells cultured as 3D organoids in monoculture (left panel), in co-culture with PSCs (middle panel), and PSC cells in co-culture with PDAC cells grown as 3D organoids (right panel). The optical redox ratio and pyruvate to lactate ratio were measured in cultures supplemented with and without 10mM exogenous pyruvate (Py). The measurements were performed on day 3 and 4 of culture. For optical redox ratio, at least n = 3 images were analyzed per condition and is subset of data shown in (e). Optical redox ratio is normalized to the PDAC cells without pyruvate on day 1. Mean pyruvate and lactate measured per day was used to assess pyruvate to lactate ratio. Error bars represent the standard deviation.

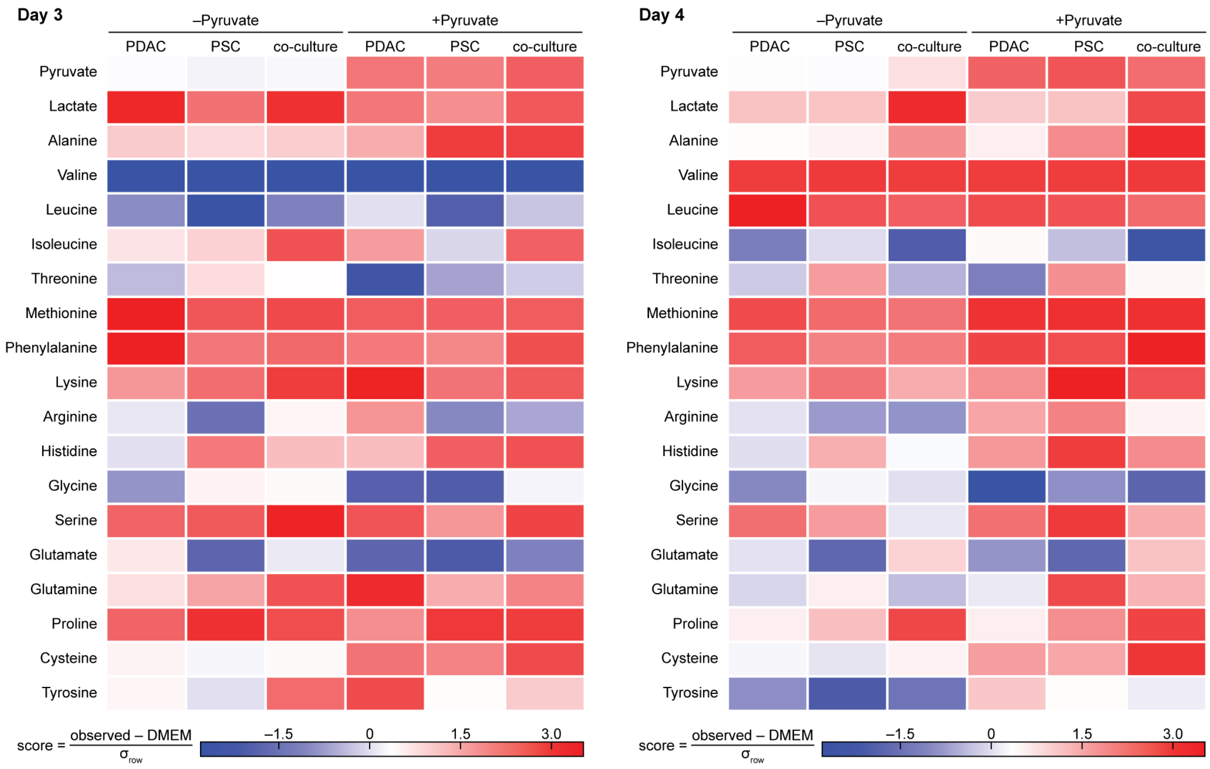


**Supplementary Figure 5.** Difference in metabolite levels measured by GC-MS in conditioned media from the indicated cells in culture compared to what is found in fresh culture media. Culture conditions evaluated include PDAC cells cultured as 3D organoids (PDAC), PSCs cultured in 3D (PSC), or PDAC cells cultured as 3D organoids in co-culture with PSCs (co-culture). Media was analyzed after 3 days of culture (left) or 4 days of culture (right) in DMEM-based media supplemented with or without (+/-) 10mM pyruvate as indicated. The score presented in the heatmap for each condition was generated by calculating the difference between each metabolite measured in conditioned media (observed) and that measured in fresh DMEM media alone (DMEM), normalized to the respective standard deviation of the metabolite levels measured for all conditions, i.e., respective metabolite row ($\sigma_{row}$). The raw data is provided in Supplementary Table 3.

**Supplementary Table 1.** Number of cells segmented for data shown in Figure (3e)

| Day | Cell type | Culture | Pyruvate | n = dishes | images per dish | n = number of cells |
|---|---|---|---|---|---|---|
| 1 | PDAC cells | monoculture | - | 3 | 2 | 172 |
| | | | + | 3 | 2 | 197 |
| | | co-culture | - | 3 | 2 | 246 |
| | | | + | 3 | 2 | 380 |
| | PSCs | monoculture | - | 3 | 2 | 57 |
| | | | + | 3 | 2 | 59 |
| | | co-culture | - | 3 | 2 | 48 |
| | | | + | 3 | 2 | 71 |
| 2 | PDAC cells | monoculture | - | 3 | 2 | 162 |
| | | | + | 3 | 2 | 180 |
| | | co-culture | - | 3 | 2 | 443 |
| | | | + | 3 | 2 | 352 |
| | PSCs | monoculture | - | 3 | 2 | 63 |
| | | | + | 3 | 2 | 158 |
| | | co-culture | - | 3 | 2 | 78 |
| | | | + | 3 | 2 | 96 |
| 3 | PDAC cells | monoculture | - | 3 | 2 | 262 |
| | | | + | 3 | 2 | 228 |
| | | co-culture | - | 3 | 2 | 414 |
| | | | + | 3 | 2 | 445 |
| | PSCs | monoculture | - | 3 | 2 | 69 |
| | | | + | 3 | 2 | 129 |
| | | co-culture | - | 3 | 2 | 102 |
| | | | + | 3 | 2 | 94 |
| 4 | PDAC cells | monoculture | - | 3 | 2 | 355 |
| | | | + | 3 | 2 | 201 |
| | | co-culture | - | 3 | 2 | 347 |
| | | | + | 3 | 2 | 258 |
| | PSCs | monoculture | - | 3 | 2 | 38 |
| | | | + | 3 | 2 | 104 |
| | | co-culture | - | 3 | 2 | 46 |
| | | | + | 3 | 2 | 91 |

**Supplementary Table 2**. Number of PSCs segmented for data shown in Figure (4e)

| Day | Group | Pyruvate | n= number of cells |
|---|---|---|---|
| 1 | non-touching | - | 31 |
| | | + | 11 |
| | touching | - | 48 |
| | | + | 71 |
| 3 | non-touching | - | 21 |
| | | + | 11 |
| | touching | - | 102 |
| | | + | 94 |
| 4 | non-touching | - | 15 |
| | | + | 14 |
| | touching | - | 46 |
| | | + | 91 |

**Supplementary Table 3.** Total ion count of metabolites analyzed by GC-MS

| Day | Culture | Pyruvate | Norvaline | Pyruvate | Lactate | Alanine | Glycine | Serine | Glutamate | Glutamine | Proline | Valine |
|---|---|---|---|---|---|---|---|---|---|---|---|---|
| N.A | DMEM | - | 15084455 | 553970 | 740208 | 85201 | 9744334 | 11728719 | 531832 | 6541063 | 116544 | 15085282 |
| 3 | PDAC | - | 25928704 | 14838184 | 57509896 | 1165272 | 16381055 | 18836523 | 1822534 | 21182121 | 449258 | 25929205 |
| | PDAC | + | 28225777 | 132167936 | 39843890 | 1697383 | 16711350 | 21129323 | 842567 | 36310753 | 392329 | 28226069 |
| | PSC | - | 30304785 | 8337679 | 44081307 | 1128474 | 21224874 | 22463438 | 910852 | 29088285 | 673542 | 30305816 |
| | PSC | + | 29752079 | 131561645 | 36439368 | 3662781 | 17474396 | 19626044 | 671759 | 27908662 | 637725 | 29751580 |
| | co-culture | - | 25835047 | 12054600 | 54340725 | 1134918 | 17977933 | 21486079 | 1555297 | 30063888 | 502175 | 25835703 |
| | co-culture | + | 19261828 | 102802040 | 32756850 | 2305871 | 13184338 | 14899538 | 734556 | 20035413 | 407791 | 19265338 |
| 4 | PDAC | - | 29519435 | 23709566 | 106600392 | 2303452 | 20201055 | 22170598 | 2639685 | 25816088 | 556962 | 29518534 |
| | PDAC | + | 24493446 | 130713448 | 80782497 | 2587837 | 14258607 | 18278344 | 1386351 | 22690146 | 482876 | 24493425 |
| | PSC | - | 37660057 | 27196813 | 131944631 | 3654676 | 28659296 | 25577775 | 1278930 | 39367898 | 1324800 | 37660561 |
| | PSC | + | 26732729 | 154611673 | 97097792 | 8195615 | 18447596 | 22498863 | 912289 | 40435119 | 1418959 | 26732318 |
| | co-culture | - | 37747945 | 67417515 | 387134121 | 11285638 | 28131560 | 18736750 | 4563478 | 30460745 | 3020796 | 37746154 |
| | co-culture | + | 26849767 | 134456602 | 231696029 | 14134446 | 17483311 | 17650684 | 3433589 | 32341170 | 2198290 | 26853318 |

| Day | Culture | Pyruvate | Leucine | Isoleusine | Threonine | Methonine | Phenylalanine | Cysteine | Lysine | Arginine | Tyrosine | Histidine |
|---|---|---|---|---|---|---|---|---|---|---|---|---|
| N.A | DMEM | - | 17451639 | 24327708 | 1742332 | 3955188 | 7915757 | 38187 | 6541530 | 1748833 | 6635237 | 3162149 |
| 3 | PDAC | - | 28065285 | 38983917 | 4373985 | 7977883 | 16296335 | 421359 | 21161974 | 3246141 | 8140105 | 5007524 |
| | PDAC | + | 31350199 | 44227611 | 3545756 | 7348326 | 14087672 | 1182719 | 36501917 | 4744240 | 12003306 | 10124840 |
| | PSC | - | 31265899 | 46082955 | 5961841 | 7981287 | 15233064 | 343769 | 29239465 | 2661475 | 8654133 | 14436643 |
| | PSC | + | 31517684 | 42979630 | 4822439 | 7655114 | 14379617 | 1157178 | 27964363 | 2841764 | 9222635 | 15586769 |
| | co-culture | - | 27882673 | 42257402 | 4858728 | 7001965 | 13370095 | 385378 | 30068021 | 3533054 | 10381698 | 9283795 |
| | co-culture | + | 21223872 | 31227564 | 3331613 | 4992181 | 10542146 | 984187 | 20027536 | 2009568 | 6540074 | 10632281 |
| 4 | PDAC | - | 33086128 | 43619997 | 5238090 | 7782327 | 14915066 | 530172 | 25787038 | 3059186 | 10038087 | 6959363 |
| | PDAC | + | 25620083 | 37101841 | 3884501 | 7000963 | 13204257 | 1775342 | 22635984 | 3205926 | 10387066 | 14352934 |
| | PSC | - | 39204020 | 56592459 | 8193553 | 8981939 | 17021007 | 369519 | 39520772 | 3322879 | 11154428 | 19974571 |
| | PSC | + | 27790964 | 39964515 | 5909612 | 7713833 | 14072061 | 1818130 | 40424538 | 3712538 | 10655947 | 22681252 |
| | co-culture | - | 38650364 | 55171686 | 6471865 | 8724924 | 17314609 | 1193473 | 30472913 | 3258198 | 12304868 | 11507648 |
| | co-culture | + | 27185497 | 38903167 | 5204657 | 7748631 | 16508145 | 3504563 | 32313410 | 3059359 | 10349201 | 16686617 |
